# Mapping sex-specific hormone–metabolite coupling in the adolescent brain: a longitudinal whole-brain spectroscopic imaging study

**DOI:** 10.64898/2026.09.07.749851

**Authors:** Edgar Céléreau, Federico Lucchetti, Zoé Schilliger, Yasser Alemán-Gómez, Patric Hagmann, Philippe Conus, Arnaud Merglen, Camille Piguet, Antoine Klauser, Daniella Dwir, Paul Klauser

## Abstract

Adolescence is marked by coordinated endocrine and brain maturation, yet how blood circulating steroid hormones relate to neurochemical change in the brain remains largely unknown. We combined longitudinal data from a novel fast whole-brain, high-resolution three-dimensional proton magnetic resonance spectroscopic imaging technique with repeated measurements of sexual hormones and adrenal steroids in the serum of 42 healthy adolescents (13–15 years; 24 females) totalizing 100 scan-visits. Longitudinal voxel-wise models separated within-individual changes from stable between-individual differences and controlled the false discovery rate across whole-brain tests. In the whole sample, increasing age was associated with higher N-acetylaspartate plus N-acetylaspartylglutamate within individuals, whereas age was positively associated with higher glutamate plus glutamine between individuals. We then observed a hormone–metabolite coupling that differed by sex and steroids. In males, within-individual increases in testosterone tracked frontal increases in glutamate plus glutamine and disseminated increases in total N-acetylaspartate. In females, higher mean estradiol between individuals was associated with higher frontal choline-containing compounds. Within-individual changes in cortisone were associated with myo-inositol and choline-containing compounds in a widespread sex-interaction effect, with positive coupling in males and negative in females. The cortisone/cortisol ratio showed a similar sex-interaction for choline-containing compounds. These findings reveal spatially distributed, sex-dependent links between steroid maturation and adolescent brain neurochemical composition and underscore the importance of differentiating within- and between-individual associations. These observational data extend predominantly structural descriptions of pubertal brain development by identifying distinct coupling of gonadal hormones with neuronal-metabolic markers and glucocorticoid interconversion with glia-weighted metabolites.

## Introduction

Adolescence is a period of brain maturation that follows puberty and coincides with important hormonal changes. It is also a window of heightened vulnerability for mental disorders with a peak incidence at 14.5 years [1]. Critically, sex-differences in mental health risk also emerge during this developmental window. For example, adolescent females become at greater risk of depression than males despite equal prevalence before puberty and this shift is concentrated during adolescence [2]. Because the gonadal-steroid environment diverges between the sexes at puberty, its effects on brain maturation are expected to be partly sex-dependent. Indeed, adolescent brain maturation shows well-documented sex differences in the timing and regional pattern of grey- and white-matter change [3,4], and pubertal status accounts for part of this divergence [5]. This brings us back to the clinical question, as sex differences in adolescent depression track pubertal status and timing more closely than chronological age [6,7].

Although these epidemiological and neurodevelopmental divergences do not establish any specific mechanism, their shared timing motivates testing whether endocrine maturation and its associations with the brain may differ by sex and could help address this question. Biologically, this period is defined by the reawakening of steroid-hormone production along several partly independent endocrine axes. Adrenarche, which precedes the outward signs of puberty, drives a pronounced rise in adrenal androgens, most notably dehydroepiandrosterone and its sulfate ester (DHEA-S), reflecting maturation of the adrenal zona reticularis [8]. Gonadarche follows, through the maturation of the hypothalamic–pituitary–gonadal axis that raises gonadal steroid output (e.g., estradiol and testosterone). Importantly, these hormones increase with age in both sexes but with different magnitude and rate, most markedly for testosterone, alongside the earlier pubertal timing of females and the greater pubertal tempo of males in mid-adolescence [9]. In parallel, the hypothalamic–pituitary–adrenal axis reorganizes itself: circulating glucocorticoid output (cortisol and cortisone) changes across puberty, and the balance of cortisol-cortisone interconversion (cortisol being regenerated by 11β-hydroxysteroid dehydrogenase type 1 and inactivated back by type 2) shifts with pubertal stage in a sex-divergent manner [10]. Longitudinal sampling shows that these endocrine trajectories are only partially tied to chronological age, so that adolescents of the same age can differ both in their absolute steroid levels and in how quickly those levels are changing [9,11].

Steroid hormones can cross the blood–brain barrier and act on the central nervous system through nuclear and membrane receptors expressed on both neurons and glia. Longitudinal structural-MRI studies have linked testosterone and estradiol to region-specific changes in grey-and white-matter volumes across puberty, with higher performances than age or Tanner stage in predicting brain changes [12,13]. Adrenal and gonadal androgens appear to exert partly opposing developmental actions: higher testosterone has been associated with cortical thinning in prefrontal regions in post-pubertal boys, whereas DHEA has been associated with pre-pubertal increases in cortical thickness across regions involved in cognitive control in both sexes [14]. Although less studied, progesterone was found to account for unique variance in default-mode surface area and orbito-affective cortical thickness [11]. Glucocorticoids, in turn, appear to influence the maturation of limbic and cortical circuits during adolescence, acting through corticosteroid receptors widely expressed in these regions [15]. Taken together, these mechanisms make steroid hormones strong candidates for metabolic modulation, especially in the adolescent brain.

Most existing evidence on steroid–brain relationships in adolescence comes from macro- and microstructural MRI measures that are largely blind to the underlying neurochemistry. Proton magnetic resonance spectroscopy (^1^H-MRS) complements these measures by quantifying neurochemical markers in vivo [16,17]. Total N-acetylaspartate (tNAA), composed of N-acetylaspartate (NAA) and N-acetylaspartylglutamate (NAAG), is synthesized in neuronal mitochondria with roles in energy metabolism, in neurotransmission (NAAG) and as a critical source for myelin lipid synthesis (NAA). Glutamate plus glutamine (Glx) captures the glutamate–glutamine cycle that underpins excitatory neurotransmission and neuronal–astrocytic metabolic coupling. Myo-inositol (Ins) is an osmolyte concentrated in astrocytes, often read as a glial marker and an unspecific index of glial density or activation. Choline-containing compounds (Cho) comprise mainly phosphocholine and glycerophosphocholine, precursors and breakdown products of membrane phospholipids, and so index membrane synthesis and turnover. Total creatine (tCr) reflects the creatine and phosphocreatine pool that buffers cellular energy through the creatine kinase system.

MR spectroscopy studies have already shown that brain neurochemistry changes measurably across development. Charting six metabolites across five regions from birth to 18 years, the steepest changes were found in the first months of life, with NAA, creatine and glutamate rising while Ins and Cho declined, before reaching relative plateaus during childhood [18]. From childhood to adulthood the trajectories are non-linear and metabolite-specific: Glx and GABA decline steeply through childhood before stabilizing in early adulthood, tCr and Cho increase (with a steeper slope in females), while tNAA and Ins remain comparatively stable [19]. Earlier quantitative MR spectroscopic imaging similarly described non-linear NAA-to-Cho trajectories in cortical grey matter alongside near-linear increases in white matter, consistent with progressive myelination [20]. These studies, however, rest on single voxels or a small number of pre-selected regions, and none has related the developing metabolic profile to the endocrine changes that accompany it. Furthermore, only one [19] did formally test sex differences.

A practical limit has been that conventional single-voxel and single-slice MRSI interrogate only a small, pre-selected region. Recent fast, whole-brain, high-resolution, three-dimensional MR spectroscopic imaging (3D-MRSI) removes this constraint, mapping several metabolites simultaneously at roughly 5-mm isotropic resolution across the entire brain within clinically feasible scan times [21]. We have previously shown that this approach reveals metabolic alterations in youth at risk for psychosis [22] and is sufficiently sensitive to build network-based structures that encapsulate aspects of the underlying biochemical brain organization [23]. Applied to typically developing adolescents, it provides a largely hypothesis-generating perspective of the locations in the brain where steroid hormones track neurochemical change, without restricting analysis to predefined regions.

A further limitation of most prior work is its cross-sectional design, which conflates two distinct questions: whether adolescents with higher hormone levels differ neurochemically from their peers (a between-subject contrast), and whether change in an individual’s hormone levels over time is accompanied by change in that same individual’s brain metabolites (a within-subject contrast). Repeated-measures data, analyzed with an explicit within- versus between-subject decomposition, help separate these effects, distinguishing dynamic, individual-level change from stable, trait-like differences and reducing confounding by inter-individual variability [12]. We therefore acquired repeated whole-brain 3D-MRSI, processed with a voxel-based pipeline adapted from our previous work [22], together with liquid chromatography-tandem mass-spectrometry (LC-MS) blood steroid panels for two or three visits per adolescent, aged between 13 and 15 years old, from the Mindfulteen cohort [24]. In this cohort, sex-specific interactions between hypothalamic–pituitary–adrenal function (cortisol-to-11-deoxycortisol ratio) and redox homeostasis have already been linked to internalizing symptoms and to white-matter microstructure in females specifically [25], providing a direct precedent for expecting sex-dependent relationships between steroid hormones and brain measures in these adolescents.

The overall aim of this study was to characterize how blood steroid hormones relate to brain metabolite concentrations measured with 3D-MRSI across adolescence, and whether these associations differ by sex or by sex-specific hormone profiles, in a longitudinal design. Specifically, we pursued three aims: (1) to characterize the age- and sex-related trajectories of a steroid panel (estradiol, progesterone, testosterone, DHEA-S, 11-deoxycortisol, cortisol and cortisone) and of five reliably resolved metabolites (tNAA, Glx, Ins, Cho and tCr); (2) to test associations of the adrenal steroids with each metabolite across the full sample, including hormone-by-sex interactions; (3) in sex-stratified models, to examine associations between each metabolite and testosterone in males, and estradiol and progesterone jointly in females. In every model, hormone effects were decomposed into within- and between-subject components. We hypothesized that androgen-related associations would be more evident for tNAA and Glx [14], whereas glucocorticoid-related associations would involve Ins and Cho [15]. Accordingly, only sparse studies have examined associations between MRS measures and steroid hormones, and none has combined 3D-MRSI, steroid hormones and a longitudinal design. Therefore, given the highly exploratory nature of this work, we deliberately remained agnostic as to which metabolites would be implicated and where such associations would be located.

## Methods

A visual sum-up of the study and the methods is provided as Fig.1.

**Fig 1.**
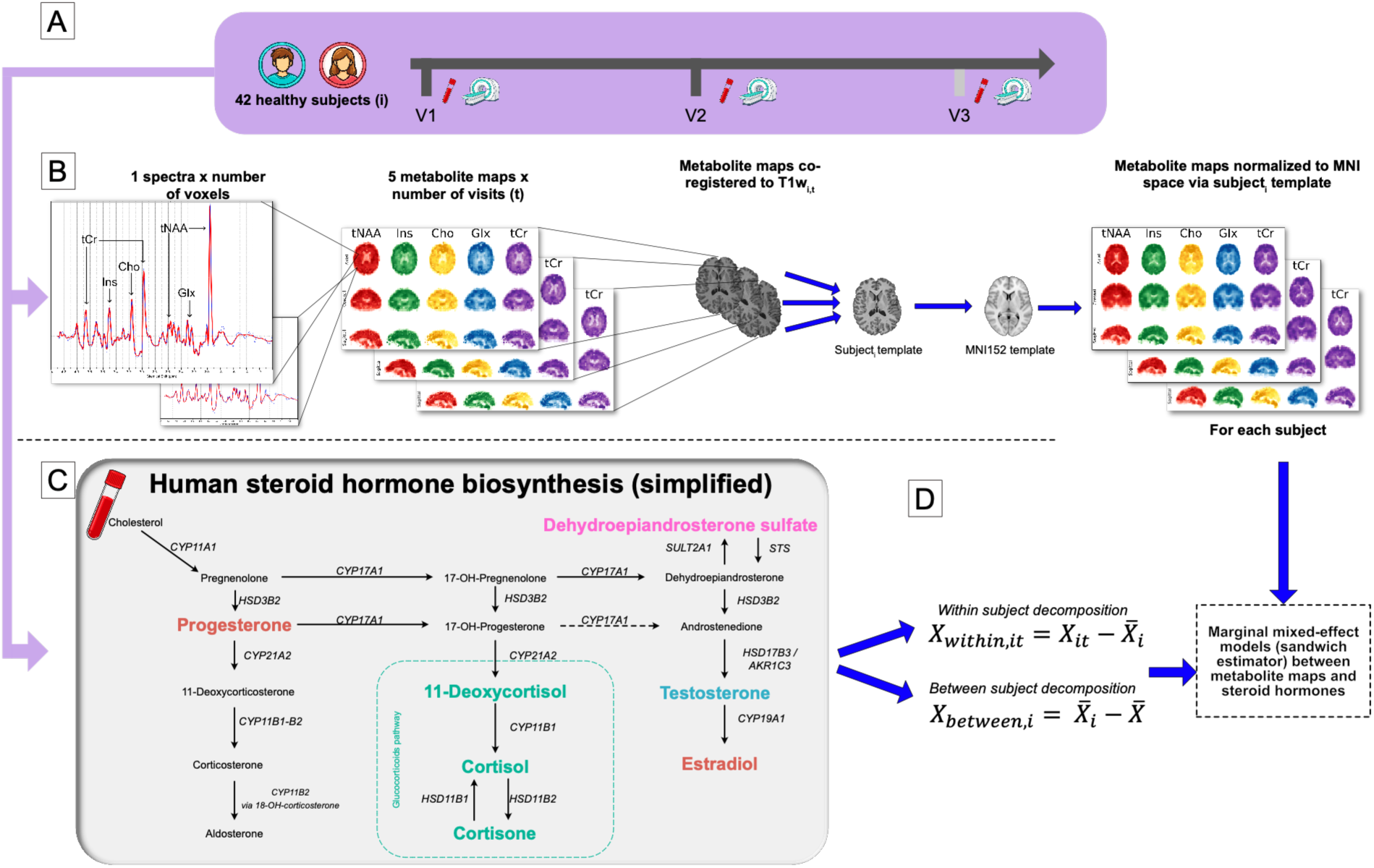
Longitudinal study design, whole-brain 3D-MRSI processing pipeline, and hormone–metabolite statistical framework. (A) Forty-two healthy adolescents (24 females, 18 males) were assessed longitudinally at up to three visits (V1, V2, V3), contributing 100 scan-visits in total. Each visit combined a fasting morning blood draw, from which serum steroids were quantified by LC-MS, with 2 days median delay whole-brain MRI and 3D-MRSI acquisition. (B) Illustration of 3D-MRSI maps specific longitudinal normalization. One spectrum per voxel is acquired across the whole brain at approximately 5-mm isotropic resolution to yield five reliably resolved metabolite maps: tNAA, Ins, Cho, Glx, tCr. For each participant, metabolite maps were co-registered to the corresponding visit’s T1w, then normalized to MNI152 standard space via an unbiased, participant-specific template built from that participant’s own T1w images across visits. (C) Simplified human steroid hormone biosynthesis pathway, from cholesterol through the glucocorticoid pathway, DHEA-S and gonadal branches with the catalyzing enzyme labeled at each step. The seven steroids quantified in this study are highlighted in color; pathway intermediates that were not measured are shown in black. (D) For each hormone, longitudinal values were decomposed into a within-subject component and a between-subject component Marginal mixed-effects models (sandwich estimator) then related the metabolite maps to these decomposed hormone components, separating dynamic within-individual coupling from stable between-individual differences.

### Participants and study design

This longitudinal study used data from the Mindfulteen project, a randomized controlled trial of a mindfulness-based intervention (MBI) in healthy adolescents from the general population. The original cohort comprised 69 participants (57% female; mean age, 14.0 ± 0.8 years) recruited between 13 and 15 years of age. Exclusion criteria were chronic somatic disease or another significant medical condition, psychotherapy during the previous 6 months, psychotropic medication during the previous month, or a DSM-IV psychiatric disorder other than an anxiety disorder or past major depressive disorder. The protocol was approved by the Ethics Committee (CCER 2018-01731); further recruitment details are provided by Piguet et al. [24]. All participants gave their written consent alongside their legal representative. Participants were randomized to receive the MBI either between V1 and V2 (early group) or after V2 (late group). The late group completed an additional post-intervention assessment, originally termed V2bis; for consistency in the present longitudinal analyses, this assessment is referred to as V3.

At each visit, body size and weight were assessed. 7 total values of height and weight were missing at visit 2 or 3 for 5 subjects. Seven missing BMI measurements at follow-up visits, corresponding to five participants, were imputed in R 4.6.1 using two-level normal model accounting for repeated observations clustered within participants implemented in the mice [26] package. Twenty imputed datasets were generated using 20 iterations and a fixed random seed of 123. Pubertal status was drawn from the K-SADS’ item assessing attainment of puberty at V1. Information regarding the use of contraceptive pills was collected but the timing of the menstrual cycle was not assessed for females.

### Serum steroid hormones

Morning blood samples were obtained after an overnight fast, and serum steroids were quantified by liquid chromatography-tandem mass spectrometry in the Clinical Chemistry Laboratory of Lausanne University Hospital. Analytes included estradiol, progesterone, testosterone, dehydroepiandrosterone sulfate (DHEA-S), 11-deoxycortisol, cortisol and cortisone. Derived indices represented cortisol synthesis/metabolism (cortisol/11-deoxycortisol and cortisone/cortisol). Ratios were calculated from the original positive concentrations before logarithmic transformation. Concentrations were expressed in nmol/L, except DHEA-S in micromol/L. One of the 100 retained scan-visits lacked the serum steroid panel; hormone analyses therefore used the resulting 99 observations without imputation. Blood samples were typically collected within 7 days before or after the scan, but 15 blood samples were collected between 7 and 21 days. Overall, the median was 2 days.

### MRI and 3D-MRSI acquisition

MR data were acquired at the Brain and Behavior Laboratory in Geneva on a 3-T Magnetom TrioTim system (Siemens Healthineers, Forchheim, Germany) with a 32-channel head coil. The T1-weighted MPRAGE acquisition used TR/TE/TI = 2300/2.98/900 ms, 1 × 1 mm in-plane resolution, 1.2-mm slice thickness, a 240 × 256 × 160 matrix, and an acquisition time of 2 min 58 s.

Whole-brain proton compressed-sensing free-induction-decay MRSI (3D 1H-CS-FID-MRSI) was acquired with TE = 1.5 ms, TR = 372 ms, flip angle = 35 degrees, field of view = 210 × 160 × 105 mm (anterior-posterior × right-left × head-foot), a 95-mm excited slab, nominal resolution = 5 × 5 × 5.3 mm3, spectral bandwidth = 2 kHz, and 512 FID points. A water-reference acquisition used TE = 1.5 ms, TR = 36 ms, flip angle = 5 degrees, resolution = 6.6 × 6.7 × 6.6 mm3, the same field of view and bandwidth, and 16 FID points. Total MRSI time was 22 min, including 2 min for the water reference. MRSinMRS [27] table is provided as Table S1.

### Reconstruction, quantification, and quality control

Measurement data were reconstructed with a low-rank model constrained by total generalized variation, incorporating simultaneous suppression of subcutaneous lipid contamination and residual water [21,28]. LCModel v6.3 [29] fitted each voxel using the unsuppressed water acquisition as reference. The fitting basis included NAA, NAAG, creatine, phosphocreatine, glycerophosphocholine, phosphocholine, myo- and scyllo-inositol, glutamate, glutamine, lactate, gamma-aminobutyric acid, glutathione, taurine, aspartate, and alanine. Because of spectral overlap, analyses used five reliably resolved maps: total N-acetylaspartate (tNAA = NAA + NAAG), total creatine (tCr = creatine + phosphocreatine), choline-containing compounds (Cho = glycerophosphocholine + phosphocholine), myo-inositol (Ins), and glutamate plus glutamine (Glx). Values were not corrected for tissue-specific T1 relaxation and are reported in institutional units.

LCModel supplied voxelwise signal-to-noise ratio (SNR), full width at half maximum (FWHM), and Cramer-Rao lower bounds (CRLB). A metabolite- and visit-specific binary quality mask retained voxels satisfying CRLB <20%, FWHM <0.1 ppm, and SNR >4. Metabolic maps were masked accordingly. Scan-level visual review assessed motion-related anatomical distortion and global cerebral coverage.

Sixty-eight participants underwent two or three 3D-MRSI acquisitions, yielding 167 acquisitions. Scan-level quality control excluded 53 acquisitions, principally because of motion-related artifacts or insufficient cerebral coverage in the metabolite quality masks, leaving 114 acceptable acquisitions. Longitudinal analyses additionally required at least two acceptable acquisitions per participant; 14 acquisitions from participants without a second usable examination were therefore not analyzed. The final longitudinal sample comprised 42 adolescents (24 female and 18 male) contributing 100 scan-visits: 37 V1, 39 V2, and 24 V3 assessments. For descriptive baseline analyses, each participant’s first retained observation was used (V1 for 37 participants and V2 for five).

### Longitudinal preprocessing

Preprocessing followed the pipeline previously described in Céléreau et al [22] and is here adapted to reduce longitudinal registration bias. Within the brain mask, high-intensity spikes exceeding the 99th percentile were identified and replaced by local 3 × 3 × 3 median inpainting. Resulting NaN and zero-valued voxels were repaired by biharmonic inpainting followed by local median filtering. Repaired voxels were smoothed with a 5-mm FWHM Gaussian kernel.

For each visit, the filtered tCr map was registered to the skull-stripped T1-weighted image using a rigid mutual-information step followed by symmetric diffeomorphic registration with Advanced Normalization Tools (ANTs v2.6.2, [30]). T1-weighted images were segmented into gray matter, white matter, and cerebrospinal-fluid probability maps with CAT12 toolbox v12.9 [31]. Tissue maps were transformed to 3D-MRSI space using the previous inverse-registration, and region-based voxelwise partial-volume correction was performed with PETPVC (v1.2.12) [32].

To avoid independently normalizing each visit, all corresponding T1-weighted images from a participant were used to construct an unbiased within-participant template with antsMultivariateTemplateConstruction2. MRSI-to-T1w-visit (from the previous tCr-to-T1w registration), T1w-visit-to-template and template-to-MNI152 transformations were finally composed and applied to each corrected metabolic map and quality mask. Registration and normalization outputs were visually checked. Normalization was done on a 5mm isotropic MNI map, corresponding to the original resolution of the 3D-MRSI maps.

### Statistical analysis

#### Demographic and hormone analyses

Non-imaging analyses were performed in R 4.6.1. At baseline, age and BMI are summarized as mean (SD), puberty as n (%), and steroid measures as median [interquartile range]. Female-male comparisons used Welch t tests for age and BMI, Fisher’s exact test for puberty, and Wilcoxon rank-sum tests for hormones.

Longitudinal hormone trajectories were assessed with linear mixed-effects models (lme4/lmerTest [33]) fitted by maximum likelihood with a participant random intercept. Positive hormone concentrations and ratios were natural log transformed. Age and BMI were decomposed into a within-participant component (visit value minus that participant’s mean) and a between-participant component (participant mean minus the population grand mean). This person-mean decomposition separated change occurring within an adolescent over time from stable differences between adolescents. Sex was sum-coded (−0.5 female, +0.5 male), so non-interaction coefficients represented the sex-averaged effect. Each model included within- and between-participant age, their interactions with sex, and within- and between-participant BMI. Benjamini-Hochberg false-discovery-rate (FDR) correction was applied to the five prespecified age/sex terms within three outcome families: sex steroids, raw adrenal steroids, and derived ratios.

#### Longitudinal voxelwise analyses

Voxelwise analyses of each of the five metabolite maps used the FSL “modified” Sandwich Estimator (SwE v1.0.3 included in FSL v6.0.7.18), which fits marginal models with robust variance estimates for longitudinal neuroimaging data [34]. For the modified SwE implementation, the subject-information file specified participant identification, visit, and a single homogeneous covariance group for all observations. For the model assessing age and sex effect, fixed effects were between-participant age, within-participant age, sex, both age-by-sex interactions. Adrenal hormones models included all participants, and each log10-transformed hormone or ratio was decomposed into between- and within-participant components; the models included both components, their interactions with sex, and between- and within-participant age and BMI. Males-only testosterone model included between- and within-participant testosterone, age, and BMI. Females-only model jointly included between- and within-participant estradiol and progesterone, age, and BMI. Planned positive and negative contrasts tested the corresponding between-participant, within-participant, and sex interaction effects when considering the whole sample. Because coverage was limited in some regions (e.g., orbitofrontal and lateral temporal cortex), analyses used a global population mask restricted to voxels with acceptable spectral quality in >90% of included acquisitions; voxels outside this mask were excluded. The ventricles and cerebellum were also excluded using the Harvard-Oxford atlas. BMI was included in all models because it influences steroid concentrations, notably through adipose aromatase activity.

Because the cohort originated from a randomized controlled trial, a secondary exploratory analysis tested whether V1-to-V2 metabolite changes differed by intervention allocation, following Piguet et al. [35]. For each metabolite, the SwE model included allocation, visit, their interaction, mean-centered baseline age, and sex. The allocation-by-visit interaction was the effect of interest. Analyses were restricted to the 34 participants with usable MRSI data at both visits.

Non-parametric inference used 5000 wild-bootstraps and threshold-free cluster enhancement (TFCE; [36]), avoiding an arbitrary cluster-forming statistic threshold for primary inference. SwE corrected-probability images were stored as 1 - p; the minimum map-level p value was therefore calculated as 1 minus the maximum voxel value. Voxelwise family-wise-error-corrected p < .05 defined a map-level effect. To address multiplicity across the planned whole-brain analyses, Benjamini-Hochberg correction (alpha = .05) was additionally applied within each metabolite and analysis stratum: 20 contrasts for all-participant hormone models, six for age/sex models, four for males, and eight for females. Primary interpretation used q <= .05; the supplementary workbook also lists maps meeting TFCE/FWE p < .05 before this across-analysis correction.

Connected components were generated only for reporting, using the suprathreshold TFCE/FWE map and 26-neighbour three-dimensional connectivity; they did not constitute a second inferential procedure. Only components larger than 10 voxels were tabulated. Peak coordinates are reported in MNI millimeters; the nearest highest-probability Harvard-Oxford cortical label was assigned, with the subcortical atlas used when no cortical label was present. Given the approximately 5-mm grid, labels are exploratory and coordinates are the primary localization.

For figures, mean values were extracted from significant components for each participant and visit. These extracted values were used descriptively and did not replace voxelwise inference.

Plots removed nuisance fixed effects while retaining the terms defining the contrast of interest and displayed fixed-effect slopes with 95% confidence intervals.

## Results

### Age and sex effect on hormones

Demographics at baseline are displayed in Table 1. The sample was comprised of 42 individuals with 57% of females. As expected, estradiol and testosterone were statistically different between males and females (p_corrected <.001). Sex differences in progesterone, 11-deoxycortisol and cortisone did not survive correction for multiple comparisons (Table 1). For gonadal steroids, sex was a significant predictor of estradiol, progesterone and testosterone levels, in both within and between-subjects (p_cor=0.002 for progesterone and p_cor<.001 for estradiol and testosterone) (Fig. 2A and 2B). However, neither the effect of age nor the sex-by-age was significant for any of the three gonadal hormones. DHEA-S showed a significant between-subjects effect of age (p_cor =0.02, Fig. 2B) but not within subjects, and no difference between sex or interaction of age by sex. Cortisol, cortisone, 11-deoxycortisol and their ratio showed no significant effect of sex, age or in sex-by-age interaction in both between and within-subjects models.

**Fig 2.**
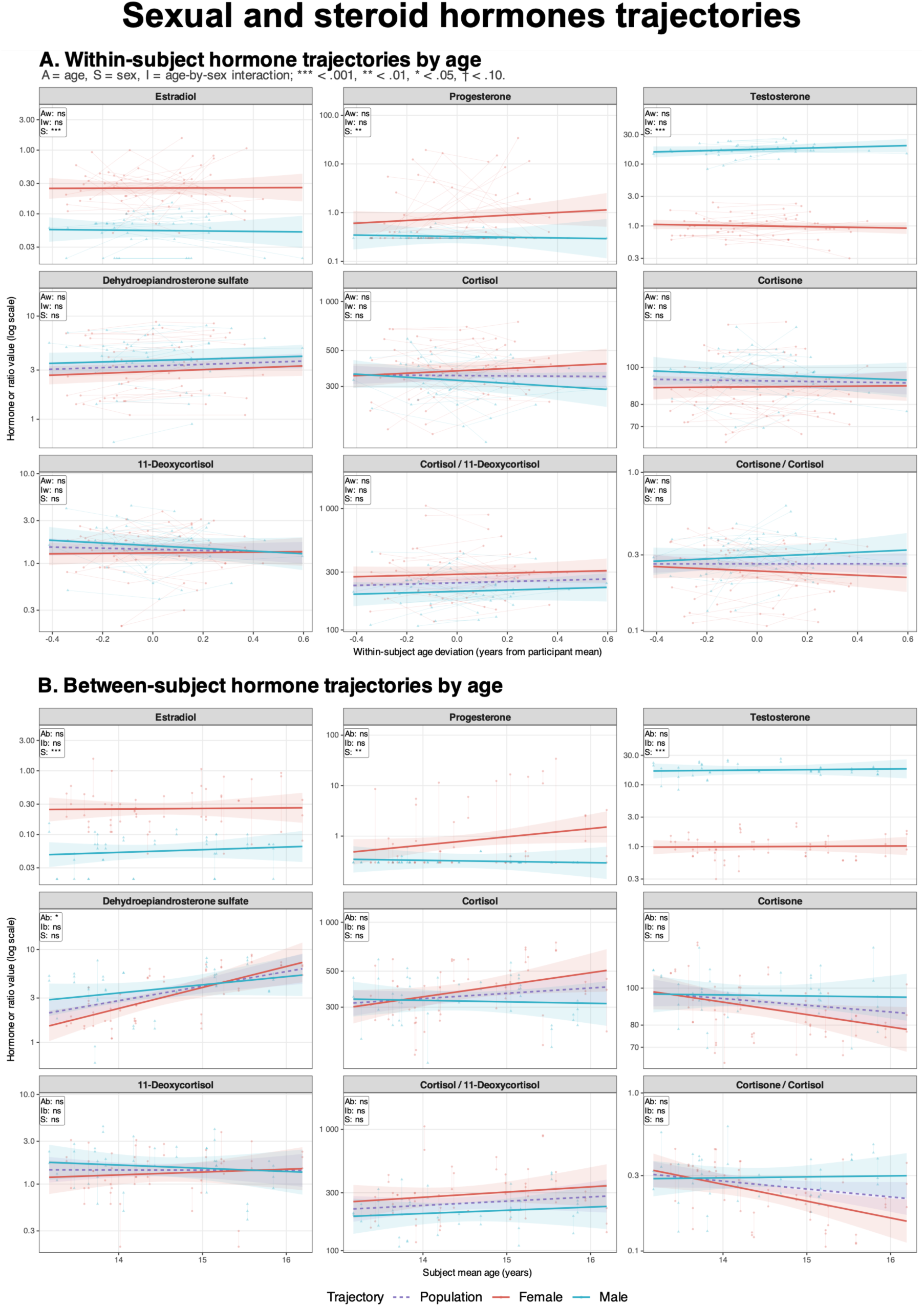
Within- and between-subject age-related trajectories of blood steroid hormones and hormone ratios. (A) Within-subject trajectories plotted against age deviation from each participant’s mean age. (B) Between-subject trajectories plotted against participants’ mean age. Points represent individual observations, with repeated measurements connected by thin lines. Females are shown as red circles and males as cyan triangles. Solid lines indicate sex-specific model estimates; the purple dashed line represents the population-level estimate and is omitted for estradiol, progesterone, and testosterone. Shaded areas indicate 95% confidence intervals. Linear mixed-effects models were fitted to natural-log-transformed outcomes and included within- and between-subject components of age and BMI, sex, interactions between sex and both age components, and a participant-specific random intercept. Annotation codes indicate the within-subject age effect (Aw), between-subject age effect (Ab), corresponding age-by-sex interactions (Iw and Ib), and sex effect (S). P values were BH adjusted within three outcome families: sex steroids, other steroid hormones, and hormone ratios. ***P < 0.001, **P < 0.01, *P < 0.05, †P < 0.10; ns, not significant.

**Table 1:**
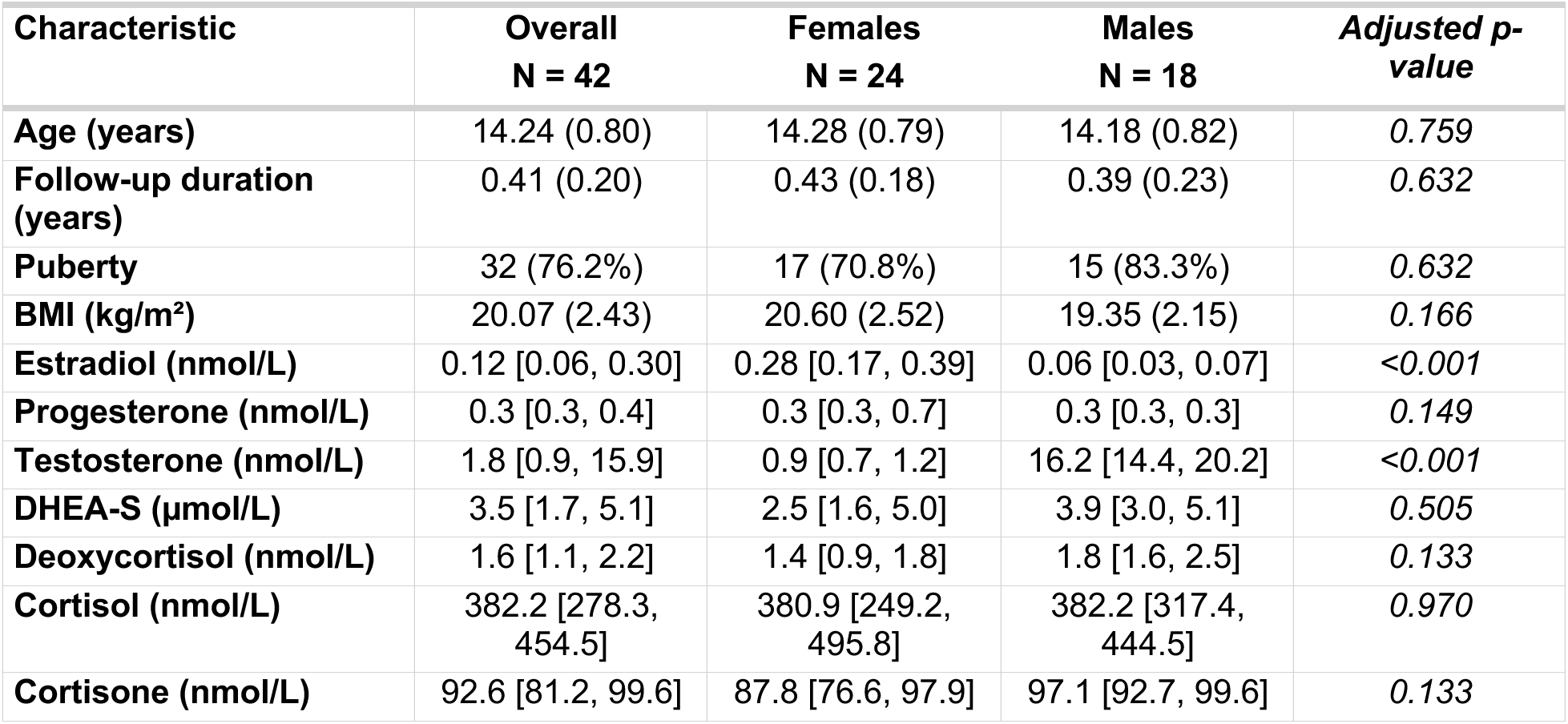
Demographics and baseline characteristics.

### Age and sex effect on brain metabolites

Given the absence of clinically measurable effects of the intervention in this cohort [35], we also first verified that the intervention had no effect on brain metabolites: mindfulness intervention yielded no significant findings for any of the five metabolites. No voxel or cluster reached significance either before (voxelwise family-wise-error-corrected p < .05) or after Benjamini–Hochberg correction for multiple comparisons.

Within subjects, age was positively associated with tNAA (p_FWE,min = .015, q_BH = .044), especially in the temporal and occipital lobes and in the corpus callosum (Fig. 3A). Between subjects, age was positively associated with Glx (p_FWE,min = .001, q_BH = .008) in a large cluster peaking and extending mostly in the frontal lobes. A pooled effect of sex was found with higher tNAA in males in widespread grey matter (p_FWE,min = .014, q_BH = .044, Fig. 3C).

**Fig 3.**
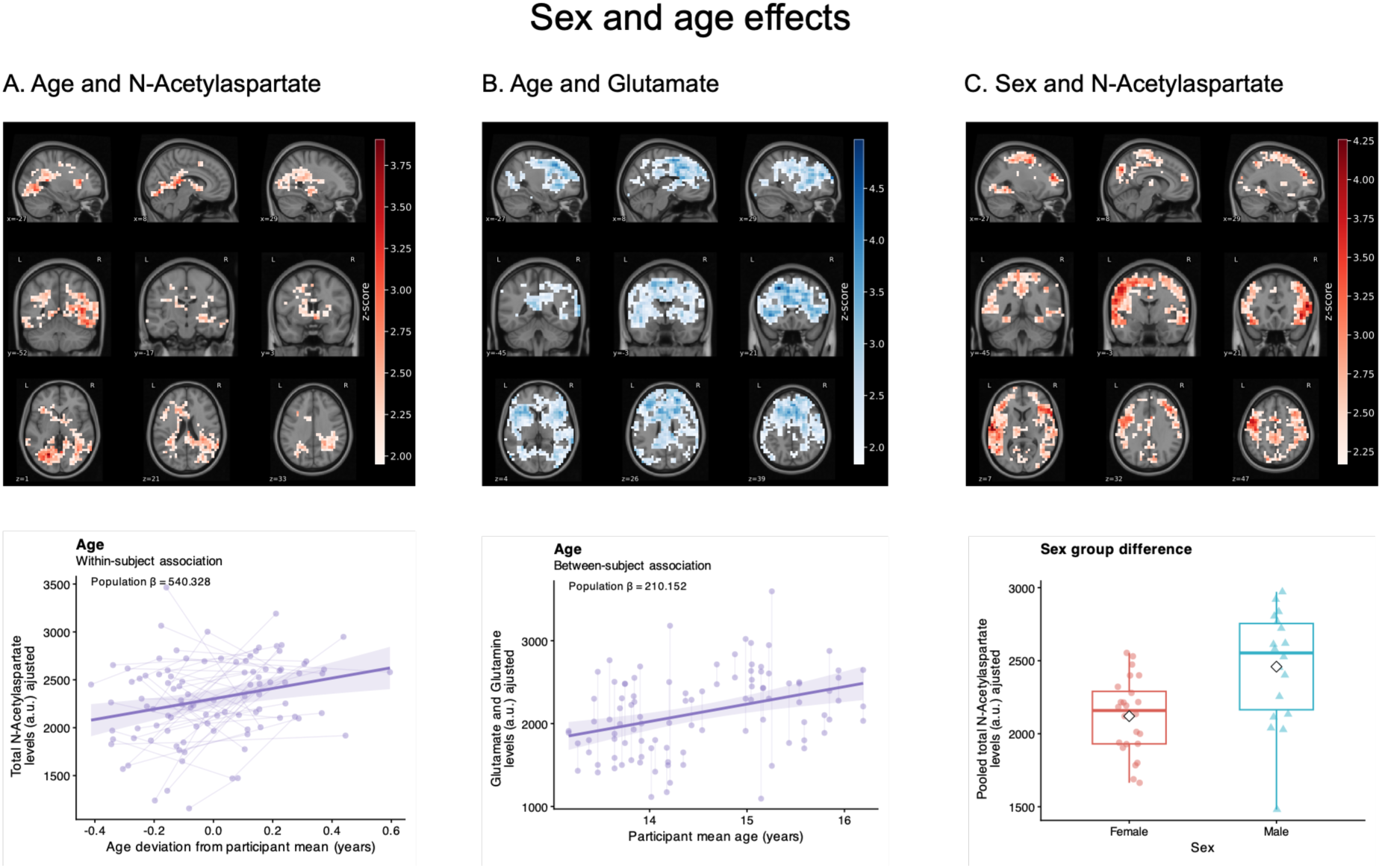
Voxel-based associations between age, sex and 3D-MRSI metabolite levels. (A) Statistical maps showing significant clusters (p < .05) in which tNAA was associated with age within subjects. Color scale represents the z-score per voxel of the marginal mixed-effect analysis. Corresponding plot below illustrates the result by using the mean metabolite signal extracted from the corresponding clusters. Points represent individual visits and faint lines connect observations from the same participant. Solid line shows fixed-effect predictions from the longitudinal model with shared areas indicating 95% confidence interval. Reported β values are unstandardized model coefficients. (B) Similarly, shows significant clusters in which Glx was associated with age between subjects. Between-subject age values are displayed on their original scale using participant geometric means. (C) Similarly, shows significant clusters in which tNAA differs by sex. Values of tNAA across visits per subjects are pooled for this specific graph.

Other findings did not meet statistical significance after correction for multiple comparisons, such as a positive association of age with tCr in occipital regions between subjects (p_FWE,min = .010, q_BH = .060) and a negative association of age with tCr in fronto-parietal regions within subjects (p_FWE,min = .023, q_BH = .068). Within subjects, age was also negatively associated with Ins, notably in the postcentral gyrus (p_FWE,min = .026, q_BH = .156).

### Sex-stratified models for the association between gonadal hormones and brain metabolites

Between female subjects, estradiol was positively associated with Cho (p_FWE,min = .005, q_BH = .040), in three clusters, the largest peaking in the frontal lobe (Fig. 4A). Within male subjects, testosterone changes were positively associated with Glx (p_FWE,min = .002, q_BH = .007, Fig. 4B) in widespread grey matter with peak p-value in frontal and temporal lobes, and with tNAA (p_FWE,min = .009, q_BH = .037, Fig. 4C) in smaller but dispersed clusters.

**Fig 4.**
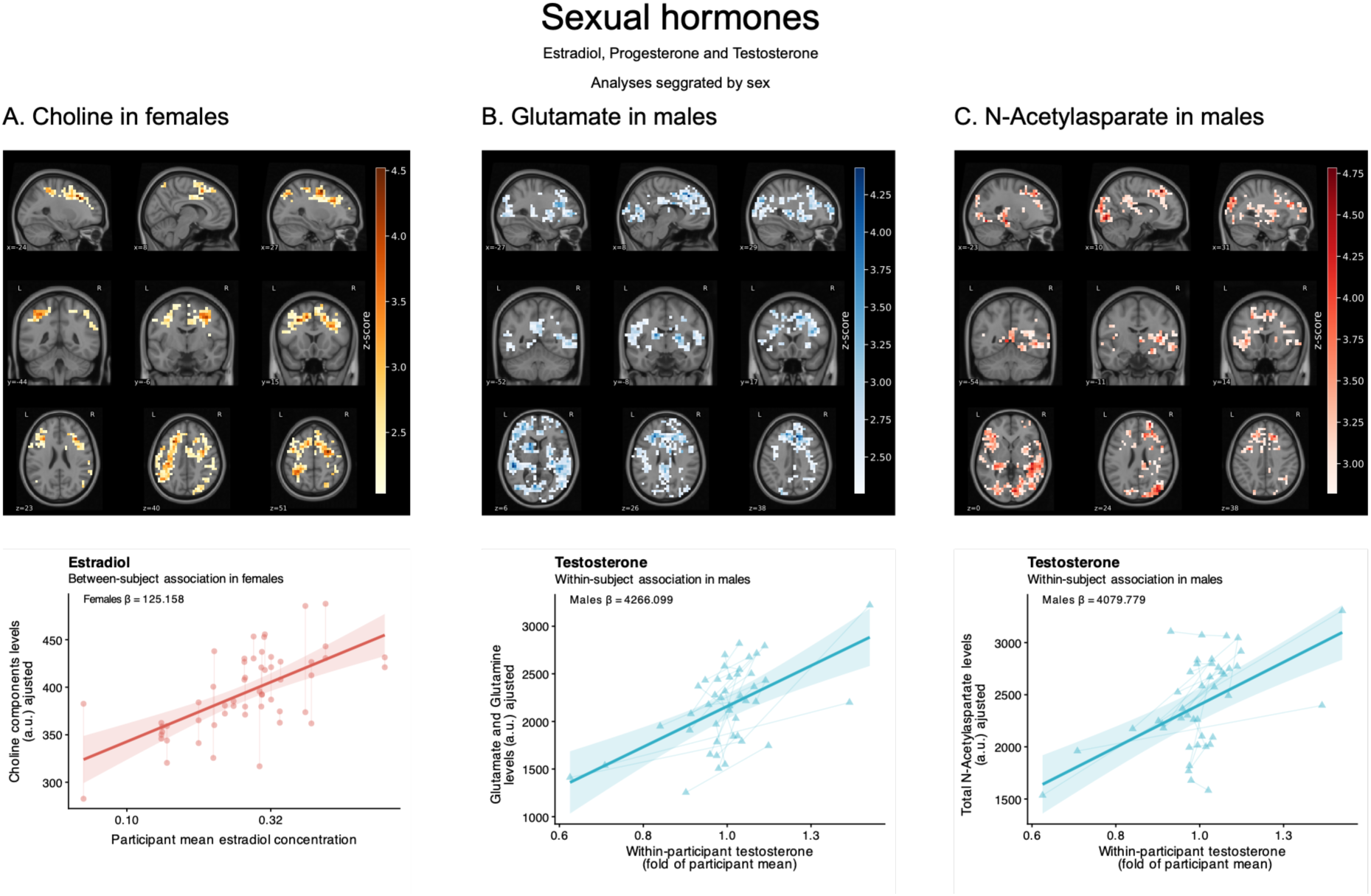
Voxel-based associations between gonadal hormones and 3D-MRSI metabolite levels, in sex-stratified models. (A) Statistical maps showing significant clusters (p < .05) in which Cho was associated with Estradiol between females. Color scale represents the z-score per voxel of the marginal mixed-effect analysis. Corresponding plot below illustrates the result by using the mean metabolite signal extracted from the corresponding clusters. Points represent individual visits and faint lines connect observations from the same participant. Solid line shows fixed-effect predictions from the longitudinal model with shared areas indicating 95% confidence interval. β corresponds to a one-unit increase on the log10 scale. Between-subject estradiol values are displayed on their original scale using participant geometric means. (B) Similarly, shows significant clusters in which Glx was associated with Testosterone within males. Within-subject values represent fold change relative to each participant’s typical hormone level. (C) Similarly to B, shows significant clusters in which tNAA was associated with Testosterone within males.

Other findings did not meet statistical significance after correction for multiple comparisons: between female subjects, positive associations emerged between estradiol and Glx (p_FWE,min = .023, q_BH = .19) in the superior parietal lobule and frontal pole, and between estradiol a nd tNAA (p_FWE,min = .017, q_BH = .139) in the postcentral gyrus (Table S2); within female subjects, a negative association emerged between both estradiol and progesterone and tCr in one small cluster for each (p_FWE,min = .013, q_BH = .107 and p_FWE,min = .038, q_BH = .131 respectively).

### Association between adrenal hormones and brain metabolites in the whole sample

A within-subject sex-interaction effect was found between cortisone and Ins (p_FWE,min = .001, q_BH = .028) in a widespread cluster encompassing almost all grey matter and parts of white matter (Fig. 5A), and Cho (p_FWE,min = .003, q_BH = .034) in a less wide cluster that still encompassed a large part of the grey matter, peaking in the postcentral gyrus (Fig. 5B); both showed a positive association in males and negative in females. A similar within-subject sex interaction between the cortisone/cortisol ratio and Cho was also retained (p_FWE,min = .003, q_BH = .034), spanning four clusters with the most important peaking in the precentral gyrus (Fig. 5C).

**Fig 5.**
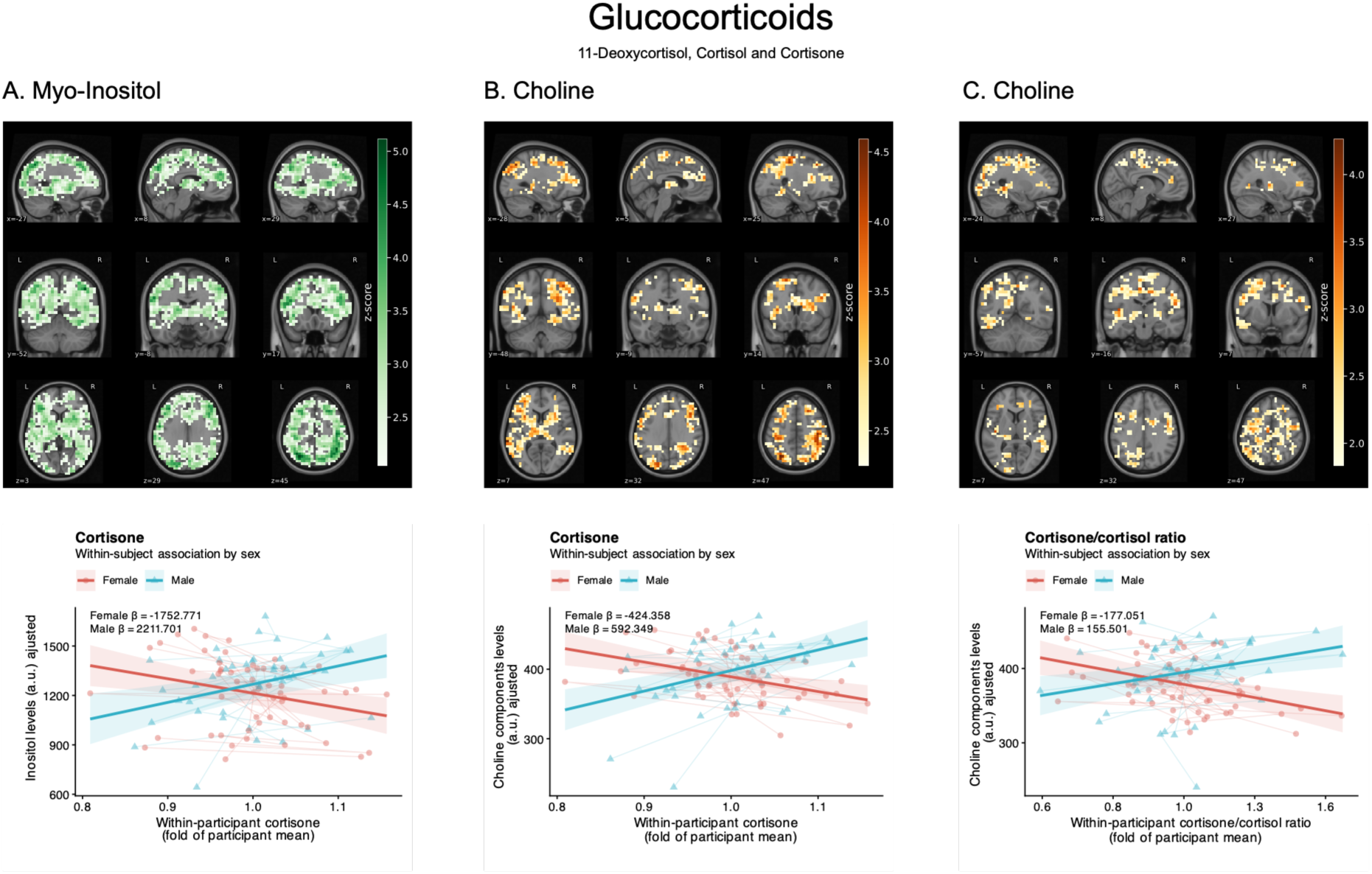
Voxel-based associations between glucocorticoid hormones and 3D-MRSI metabolite levels. (A) Statistical maps showing significant clusters (p < .05) in which the within-subject association between Cho and Cortisone differed by sex. Color scale represents the z-score per voxel of the marginal mixed-effect analysis. Corresponding plot below illustrates the result by using the mean metabolite signal extracted from the corresponding clusters. Points represent individual visits with red points for females and cyan triangles for males and faint lines connect observations from the same participant. Solid line shows fixed-effect predictions from the longitudinal model with shared areas indicating 95% confidence interval. β corresponds to a one-unit increase on the log10 scale. Within-subject values represent fold change relative to each participant’s typical hormone level. (B) Similarly, shows significant clusters in in which the within-subject association between Cho and Cortisone differed by sex. (C) Similarly, shows significant clusters in in which the within-subject association between Cho and Cortisone/cortisol ratio differed by sex.

A non-BH-significant but recurrent pattern was found for tCr and Ins: positive associations between-subject with cortisol and DHEA-S and negative with the cortisone/cortisol ratio. Several clusters were spatially extensive, including the Ins association with cortisol encompassing most white and grey matter (p_FWE,min = .008; q_BH = .118), and a similar pattern for the DHEA-S associations with tCr (p_FWE,min = .007; q_BH = .132) and Ins (p_FWE,min = .012; q_BH = .118) encompassing most posterior white matter. The negative cortisone/cortisol ratio association with Ins (p_FWE,min = .018; q_BH = .123) also encompasses a wide white-matter cluster (Table S2).

Finally, within-subject sex-interaction was also observed between cortisone and tNAA, with a positive association in widespread grey matter in males and a negative association in females (p_FWE,min = .017, q_BH = .336). Smaller clusters of the same interaction were also observed with Glx, notably in the left lateral occipital cortex and in both middle frontal gyri (Table S2).

## Discussion

In adolescents scanned two or three times between 13 and 15 years, we combined whole-brain 3D-MRSI with LC-MS steroid panels, decomposing each hormone effect into within- and between-subject components. Across this narrow window, gonadal steroids (estradiol, progesterone, testosterone) showed robust sex differences but no significant age-related change or sex-by-age divergence. DHEA-S increased with age independently of sex, and the glucocorticoid axis showed no significant effect of sex, age, or their interaction. Age was associated with higher tNAA within subjects and with higher Glx between subjects, alongside a widespread main effect of sex on tNAA (higher in males). Gonadal steroids showed an asymmetric pattern: in males, within-subject increases in testosterone tracked increases in Glx and tNAA, whereas in females the only surviving gonadal association was between-subject, linking estradiol to Cho. Glucocorticoid markers, by contrast, produced sex-dependent within-subject couplings with the glia-weighted metabolites Ins and Cho: positive in males and negative in females. These findings suggest that in this window the gonadal axis relates preferentially to neuronal-metabolic markers in males, while the adrenal axis relates to glial-metabolic markers in a sex-divergent manner.

### Age and sex effects on blood steroids

Sex differences in estradiol, progesterone and testosterone dominated both baseline and longitudinal models, consistent with divergence of gonadal output after gonadarche [8,9]. However, no significant main effect of age, nor age-by-sex interaction, was detected for any gonadal steroid. Our participants had a mean age of 14±0.8 year at baseline and a mean follow-up of 0.4 years, giving a total observation window with limited statistical power to reproduce large similar cross-sectional or longitudinal follow-up [9,37]. Moreover, between individuals, the age of steepest testosterone rise in males varies markedly [37] while estradiol and progesterone vary with the menstrual cycle in females.

DHEA-S showed a significant between-subject association with age and a positive but non-significant within-subject slope. Repeated-measures modelling of adrenal hormones in the CATS cohort showed that between-individual variance in adrenarcheal timing far exceeds variance in tempo, and that hormone level at 8–9 years predicts relative level through early adolescence [38]. By 13–15 years, most DHEA-S variance is therefore trait-like. Glucocorticoids showed no significant age or sex effect. This fits the concept of pubertal recalibration of the HPA axis, in which cortisol regulation is coupled to pubertal and adrenal hormones rather than to chronological age [39].

### Age and sex effects on brain metabolites

Within-subject increases in age were associated with higher tNAA in a cluster encompassing important parts of the posterior white matter (parieto-occipito-temporal, splenium of the corpus callosum). NAA is exported from neurons to oligodendrocytes, where aspartoacylase liberates the acetate moiety used for myelin lipid synthesis [40]; a within-individual rise concentrated in posterior white matter is therefore read as ongoing myelination, with increased mitochondrial-energetic demand as a secondary contributor. Although a decrease with age is usually found with tNAA, our finding on a short age span during adolescence could be supported with previous non-linear white-matter tNAA levels described in earlier multi-voxel or single-voxel MRS work [19,20].

The between-subject Glx increase peaked in the frontal lobe and did not overlap the posterior tNAA age clusters. Frontal cortex matures late: adolescence is proposed to be a critical period for prefrontal circuit refinement, during which excitatory–inhibitory glutamatergic circuitry and functional engagement are progressively strengthened [41]. A between-subject increase in frontal Glx among older adolescents is compatible with this protracted strengthening of glutamatergic circuitry, for which the region’s high aerobic glycolysis, as a signature of synaptic remodeling [42], offers a plausible metabolic substrate. This maturational difference appears to be more cumulative than incremental and thus appears in a between-subject difference rather than within-subject. This finding is discordant with Thomson et al [19], who report Glx declining between childhood and adulthood linearly. However, our main effect is localized quite far from the chosen voxel in this study (i.e., parietal posterior), which may explain this discrepancy.

Sex-difference in tNAA across widespread grey matter was not accompanied by an age-by-sex interaction, indicating a trait-like difference rather than divergence emerging over the observation window. We already reported a similar result in cross-sectional design using the same sample and 2 replication samples, alongside a tNAA/tCr similar widespread difference and we interpreted this difference as a sexual metabolic dimorphism [43]. Interestingly, the absence of age-by-sex interaction may suggest that this difference is already established before the window sampled here. Since we have previously observed important white matter maturation differences in these same subjects [44], a metabolic dimorphism that may underly these maturational changes could emerge before 13 years old. Of note, a comparable difference has been reported in adults with lower tNAA/tCr in parietal cortex of cisgender women compared to cisgender men [45].

Two further age effects did not survive correction but are developmentally coherent. Ins and tCr both decreased within subjects in cortical grey matter, consistent with the pronounced cortical thinning during adolescence [46], while tCr increased between subjects in white matter, most markedly frontally, in line with the protracted maturation of frontal associated white matter over this period [47]. This directional and spatial dissociation is consistent with grey-matter synaptic and glial consolidation proceeding alongside white-matter maturation, the latter carrying a higher creatine-buffered energetic cost.

### Testosterone and neuronal metabolites in males

In males, within-subject increases in testosterone were associated with increases in Glx across fronto-temporo-occipital grey matter, and with increases in tNAA in smaller clusters in the same regions. Because both age components were included in the model, these coefficients reflect testosterone deviating from its age-expected value. Consistent with this, the testosterone and age effects occupied distinct territory.

To our knowledge, no previous study has related endogenous testosterone to Glx in the developing human brain. The only prior human evidence linking testosterone to glutamatergic metabolites comes from gender-affirming hormone therapy, in which exogenous, supraphysiological testosterone was administered to adult trans men [45,48]. Spurny-Dworak et al found a reduced hippocampal GABA+/tCr after testosterone administration, an indirect excitation-inhibition shift in the same direction as our increased Glx. Androgens are also plausible modulators of excitatory synaptic architecture. In animal models, gonadectomy reduces synaptic spine density on CA1 pyramidal neurons, an effect rapidly reversed by testosterone or dihydrotestosterone [49], and androgens modulate spinogenesis through both intracellular and membrane androgen receptors [50,51].

The parallel testosterone-tNAA association connects to the higher tNAA observed in males as compared to females: a continuing androgenic influence on neuronal-metabolic markers within males would be expected to sustain, and perhaps widen, a sex difference established during development. Direct human evidence remains scarce but concordant: in men undergoing androgen-deprivation therapy, testosterone decrease mediated hippocampal NAA/Cr fell over six months, and this was associated with cognitive decline [52]. Altogether, this points toward an important link between brain tNAA and testosterone in the male population, although it is important to note that this association seems to be only linked relatively to a subject’s own testosterone level, and not the absolute concentration of circulating testosterone.

### Estradiol and progesterone in females

Among females, the only gonadal association surviving correction was between-subject: higher mean estradiol was associated with higher Cho in three clusters, the largest peaking in the middle frontal gyrus bilaterally. Sub-threshold between-subject associations recurred for Glx and tNAA, and small negative within-subject associations of estradiol and progesterone with tCr were observed.

Cho index membrane phospholipid synthesis and turnover, including myelin-related processes, which may be modulated by estrogen [53]. A steepest increase in Cho in females was also observed in a previous MRS study [19]. Beyond myelin-related turnover, elevated Cho could plausibly also reflect the membrane remodeling that accompanies synaptic pruning, which begins at puberty in human cortex and proceeds over a protracted period [54]. As females reach peak grey-matter volume one to two years earlier than males [55], a greater contribution of pruning-related membrane turnover may occur in females, though our data cannot establish this link directly and the Cho signal is not specific to any particular membrane process.

Two considerations bear the absence of a significant within-subject effect in females. First, menstrual cycle phase was not recorded, and only 2 to 3 measures of estradiol and progesterone per participant does not allow to make any assumption on this phase; therefore, the between-subject component probably captures a less noisy effect. Second, females in this cohort are on average more advanced through puberty, and much estradiol-related remodeling may predate 13 years. Multimodal diffusion analysis in the same cohort indicated more advanced white-matter maturation in females, linked to sexual hormone, HPA-axis and redox homeostasis [44]. The nominally significant associations between tNAA and Glx with between-females estradiol localized predominantly in the frontal lobe could also be consistent with this hypothesis, in contrast to the more dispersed increase observed in males with testosterone.

However, a within-subject effect appeared with a positive association between estradiol and tCr in the left para-hippocampal zone, and a negative association between progesterone and tCr in a very small nearby cluster. Although they did not survive correction for multiple comparisons, we can speculate that our sample was underpowered to detect such an effect, which is in line with a previous publication that found an asymmetric fluctuation of tCr in females during the menstrual cycle [56].

### Glucocorticoid metabolism and sex-divergent glial trajectories

The clearest sex-dependent findings arose in the adrenal models. Within-subject changes in cortisone showed a sex interaction with Ins across almost all grey matter and part of the white matter, and with Cho over a large grey-matter cluster peaking in the postcentral gyrus; the cortisone/cortisol ratio showed the same interaction with Cho. In each case the association was positive in males and negative in females, with a same-signed non-surviving interaction for tNAA and, in smaller clusters, Glx.

That these effects involve cortisone and the cortisone/cortisol ratio rather than cortisol points to a potential involvement of the 11β-hydroxysteroid dehydrogenase type 1 (11β-HSD1), which regenerates cortisol intracellularly from cortisone and amplifies local glucocorticoid action independently of circulating levels. The enzyme is expressed across the human brain, including widespread neocortex [57], and its inhibition blocks stress-induced suppression of hippocampal synaptic potentiation, showing that local rather than systemic glucocorticoid metabolism governs functional outcome [58]. Crucially, brain 11β-HSD1 is expressed in astrocytes and microglia, where it regulates glial reactivity and neuroinflammatory state [59,60], which may be reflected by Ins, an astrocytic osmolyte and marker of glial activation [61], and by Cho, a marker of membrane turnover. The cortisone/cortisol ratio is thus better read as an index of glucocorticoid interconversion capacity than of exposure, and this interconversion, not absolute cortisol, tracked glial-metabolic change in this age range. It is also important to note that Ins and Cho were already reported being particularly collinear in adolescence, compared to childhood and adulthood [19].

Adolescence is a plausible, and potentially sensitive, window for such effects: cortical glucocorticoid-receptor expression is itself being reorganized, shifting from astrocytic toward pyramidal-neurons localization across development [62], while glucocorticoids continue to shape dendritic architecture and plasticity [15]. This could also help understand the non-significant, but similar effect of cortisone on tNAA and Glx, this progressive shift being progressively apparent but with a lower magnitude at this age compared to glial markers.

We interpret the sex interaction as males and females occupying opposite phases of a shared glial trajectory rather than differing qualitatively in glucocorticoid response. If females have largely completed the expansion phase of glial and myelin-related maturation, maturing glucocorticoid metabolism would coincide with consolidation, expressed as a negative coupling between rising cortisone and Ins/Cho, whereas still-expanding males show the reverse. This interpretation is consistent with the more advanced female white-matter maturation and sex-specific HPA-axis–redox interactions reported in this cohort [25,44]. Such divergence is mechanistically plausible given that 11β-HSD1 is under sex-specific gonadal-steroid control [63].

A second, non-exclusive possibility is that the sexes differ not in phase but in the direction of their glial response to glucocorticoids, such that a comparable subject-specific rise in local glucocorticoid availability promotes glial metabolism in males and represses it in females. Sex-dependent effects of stress and glucocorticoids on cortical glia have been documented in rodents: chronic stress differentially engages astrocytes and microglia at glutamatergic synapses in a sex-and development-dependent manner [64], within a broader pattern of sex differences in stress-response systems [65]. This intra-individual sex-specific coupling between glucocorticoid signaling and glia could accumulate across adolescence and would contribute to widening the sex gap apparent by early adulthood.

The between-subject adrenal associations, though not surviving correction, were internally coherent: tCr and Ins were positively associated with cortisol and DHEA-S and negatively with the cortisone/cortisol ratio, over spatially extensive clusters. Because the ratio is inversely related to active glucocorticoid availability, these signs point in the same direction, and a cortisol–myo-inositol relationship has been reported previously [66]. Moreover, it supports the hypothesis that these adrenal steroids are related to glial metabolism.

### Strength and limitations

To our knowledge, this is the first study to apply whole-brain, high-resolution 3D-MRSI in a longitudinal design to healthy adolescents, and the first to analyze repeated whole-brain spectroscopic imaging voxelwise rather than through single voxels or pre-selected regions. Earlier developmental spectroscopy has relied on single voxels or region-of-interest averages [18–20,67], which limits anatomical specificity and cannot localize where metabolic change occurs; our unbiased within-participant registration and voxelwise inference recover this spatial information across the brain.

A second strength concerns the breakdown of hormone levels in within- and between-subject component, which is well established [68] and has been applied to hormone–brain relationships in adolescence [12,39], but to our knowledge this is the first time these hormone effects have been separated in whole-brain spectroscopic mapping. The two components answer genuinely different questions: whether an individual’s metabolite levels change as that individual’s hormones change, and whether adolescents differing in hormone levels differ metabolically. Here testosterone and glucocorticoid interactions acted within subjects and estradiol between subjects, a structure that a single pooled coefficient would have masked. Because most endocrine–brain evidence in adolescence reports only the between-subject contrast, this bears directly on how such associations are interpreted and compared and argues in favor of repeated-measures designs in developmental neuroendocrine imaging.

Several limitations temper these findings. The sample is modest (42 adolescents, 100 scan-visits) and the age range narrow, limiting power for age effects and precluding modelling of non-linear trajectories. Whole-brain coverage was incomplete in the orbitofrontal and lateral temporal cortices. This is a well-recognized constraint of MRSI arising from their proximity to air-filled paranasal cavities and the resulting magnetic-field inhomogeneity. Accordingly, the absence of effects in these regions is uninformative. Blood sampling and brain MRI were not exactly simultaneous (median delay of 2 days), menstrual cycle phase was not recorded, and single morning serum measurements do not capture diurnal or pulsatile HPA dynamics. Finally, the design is observational: because both the gonadal and adrenal axes are under central hypothalamic–pituitary control, the associations reported here should not be read as unidirectional hormonal effects on brain tissue.

## Conclusion

Using repeated whole-brain 3D-MRSI and LC-MS steroid panels in adolescents aged 13–15, testosterone was coupled to within-individual increases in neuronal-metabolic markers in males, estradiol was related to choline as a stable between-individual difference in females, and glucocorticoid interconversion markers were coupled to glia-weighted metabolites in opposite directions in the two sexes. Sex differences in adolescent brain maturation have so far been characterized mostly through structural imaging; our results add a spatially resolved neurochemical dimension to that picture and locate the clearest sex divergence in the coupling between glucocorticoid metabolism and glia-weighted metabolites. This raises the possibility that the sexes differ not only in the timing of maturation but in how their glia respond to glucocorticoid signaling, a difference that only a whole-brain metabolic approach can resolve. This study therefore advocates for the large inclusion of 3D-MRSI in neurodevelopmental studies.

## Acknowledgments

We gratefully acknowledge the entire Mindfulteen team who contributed to the acquisition of the 3D-MRSI dataset for Geneva 3T, along with all participants.

