## Supplementary Material for "Mapping sex-specific hormone–metabolite coupling in the adolescent brain: a longitudinal whole-brain spectroscopic imaging study"

#### Table S1: MRSinMRS information

| **Category** | **Mindfulteen 3D-MRSI protocol** |
|---|---|
| Scanner | 3T Magnetom TrioTim (Siemens) |
| RF coils | 32Rx ch 1H head coil |
| Coil elements | all (32) |
| Sequence | 3D 1H-FID-MRSI (CS-accelerated) |
| Orientation | Transverse |
| Rotation (deg) | -2 |
| TE (ms) | 1.5 |
| TR (ms) | 372 |
| Averages | 1 |
| Flip angle (°) | 35 |
| FOV (mm) | 210 × 160 × 105 |
| Slab thickness (mm) | 95 |
| Slabs | 1 |
| Resolution ($mm^3$) | 5 × 5 × 5.3 |
| Spectral bandwidth (Hz) | 2000 |
| FID points / Vector size | 512 |
| Acquisition duration (ms) | 256 |
| Matrix size | 42 × 32 × 20 |
| Water reference TE (ms) | 1.5 |
| Water reference TR (ms) | 36 |
| Water reference flip angle (°) | 3 |

| | |
|---|---|
| Water reference resolution ($mm^3$) | 6.6 × 6.7 × 6.6 |
| Water reference FID points | 16 |
| Averaging mode | N.A. |
| Water suppr. | WET water suppr. |
| Water suppr. BW (Hz) | 60 |
| Spectral suppr. | None |
| Measurements | 1 |
| Saturation bands | 2 bands, 20 mm thickness |
| Compressed sensing | Acceleration factor 3.3 |
| Preparation scans | 4 |
| Dimension | 3D |
| Delta frequency (ppm) | 0.00 |
| Phase encoding | Elliptical |
| Remove oversampling | N.A. |
| Shim mode | Advanced |
| Data processing | Low-rank + TGV reconstruction; lipid/water removal |
| Quantification | LCModel |
| Metabolite basis set (LCModel) | NAA, NAAG, Cr, PCr, GPC, PCh, mI, sI, Glu, Gln, Lac, GABA, GSH, Tau, Asp, Ala |
| Combined metabolites | tNAA (NAA+NAAG), tCr (Cr+PCr), Cho (GPC+PCh), Ins (mI), Glx (Glu+Gln) |
| Quality metrics | SNR, CRLB, FWHM |

### Table S2: Supplementary voxel-based analysis results

This table reports map-level TFCE/FWE p < .05 clusters with >10 voxels; anatomical labels are exploratory peak lookups.

**31 significant analyses • 62 clusters >10 voxels • 30 analyses represented • 9 analyses retained after BH**

| Population | Metabolite | Hormone / predictor | Effect | Cluster | k (voxels) | Volume (mm³) | Hemisphere | Atlas peak label | Atlas prob. | Peak x (MNI mm) | Peak y (MNI mm) | Peak z (MNI mm) | Cluster peak p-FWE | Analysis min p-FWE | BH q-value | BH status |
|---|---|---|---|---|---|---|---|---|---|---|---|---|---|---|---|---|
| All participants | Choline | Cortisol / 11-deoxycortisol ratio | Sex interaction — between-subject component | 1 | 62 | 8,144 | Left | Cingulate Gyrus, anterior division | 9% | -11.6 | 33.8 | 14.4 | 0.0302 | 0.0302 | 0.2013 | Not retained after BH |
| All participants | Choline | Cortisone | Sex interaction — within-subject component | 1 | 2,423 | 318,272 | Right | Postcentral Gyrus | 25% | 34.2 | -27.2 | 50.0 | 0.0034 | 0.0034 | 0.0340 | Retained after BH |
| All participants | Choline | Cortisone / Cortisol ratio | Sex interaction — within-subject component | 1 | 1,868 | 245,371 | Right | Precentral Gyrus | 28% | 49.4 | -1.8 | 34.8 | 0.0034 | 0.0034 | 0.0340 | Retained after BH |
| All participants | Choline | Cortisone / Cortisol ratio | Sex interaction — within-subject component | 2 | 37 | 4,860 | Left | Frontal Operculum Cortex | 23% | -37.0 | 28.7 | 4.3 | 0.0304 | 0.0034 | 0.0340 | Retained after BH |
| All participants | Choline | Cortisone / Cortisol ratio | Sex interaction — within-subject component | 3 | 14 | 1,839 | Right | Cingulate Gyrus, anterior division | 89% | 3.7 | 38.8 | 14.4 | 0.0380 | 0.0034 | 0.0340 | Retained after BH |
| All participants | Choline | Cortisone / Cortisol ratio | Sex interaction — within-subject component | 4 | 13 | 1,708 | Right | Lingual Gyrus | 29% | 24.0 | -62.8 | -0.8 | 0.0364 | 0.0034 | 0.0340 | Retained after BH |
| All participants | Creatine | Cortisol | Between-subject effect | 1 | 1,363 | 179,036 | Left | Superior Frontal Gyrus | 48% | -21.7 | 28.7 | 50.0 | 0.0174 | 0.0174 | 0.1740 | Not retained after BH |
| All participants | Creatine | Cortisol | Between-subject effect | 2 | 85 | 11,165 | Right | Occipital Fusiform Gyrus | 15% | 34.2 | -62.8 | -5.9 | 0.0378 | 0.0174 | 0.1740 | Not retained after BH |
| All participants | Creatine | Cortisol | Between-subject effect | 3 | 27 | 3,547 | Right | Precuneous Cortex | 59% | 8.8 | -52.7 | 9.3 | 0.0412 | 0.0174 | 0.1740 | Not retained after BH |
| All participants | Creatine | Cortisol | Between-subject effect | 4 | 13 | 1,708 | Left | Lingual Gyrus | 61% | -16.7 | -52.7 | -5.9 | 0.0454 | 0.0174 | 0.1740 | Not retained after BH |
| All participants | Creatine | Cortisone | Sex interaction — within-subject component | 1 | 19 | 2,496 | Right | Frontal Pole | 26% | 44.3 | 33.8 | 24.6 | 0.0320 | 0.0320 | 0.6399 | Not retained after BH |
| All participants | Creatine | Cortisone / Cortisol ratio | Between-subject effect | 1 | 198 | 26,008 | Left | Superior Frontal Gyrus | 34% | -11.6 | 33.8 | 50.0 | 0.0304 | 0.0304 | 0.2026 | Not retained after BH |
| All participants | Creatine | Cortisone / Cortisol ratio | Between-subject effect | 2 | 20 | 2,627 | Left | Precentral Gyrus | 34% | -6.5 | -32.3 | 60.2 | 0.0462 | 0.0304 | 0.2026 | Not retained after BH |
| All participants | Creatine | Cortisone / Cortisol ratio | Between-subject effect | 3 | 20 | 2,627 | Right | Superior Frontal Gyrus | 16% | 8.8 | 43.9 | 39.8 | 0.0426 | 0.0304 | 0.2026 | Not retained after BH |
| All participants | Creatine | Cortisone / Cortisol ratio | Between-subject effect | 4 | 11 | 1,445 | Right | Juxtapositional Lobule Cortex (formerly Supplementary Motor Cortex) | 28% | 8.8 | -12.0 | 65.3 | 0.0472 | 0.0304 | 0.2026 | Not retained after BH |
| All participants | Creatine | DHEA-S | Between-subject effect | 1 | 2,636 | 346,251 | Right | Postcentral Gyrus | 21% | 13.8 | -37.4 | 60.2 | 0.0066 | 0.0066 | 0.1320 | Not retained after BH |
| All participants | Glx | Cortisone | Sex interaction — within-subject component | 1 | 202 | 26,534 | Left | Lateral Occipital Cortex, superior division | 64% | -31.9 | -67.9 | 44.9 | 0.0178 | 0.0178 | 0.3559 | Not retained after BH |
| All participants | Glx | Cortisone | Sex interaction — within-subject component | 2 | 127 | 16,682 | Left | Middle Frontal Gyrus | 45% | -31.9 | 13.4 | 55.1 | 0.0312 | 0.0178 | 0.3559 | Not retained after BH |
| All participants | Glx | Cortisone | Sex interaction — within-subject component | 3 | 124 | 16,288 | Right | Middle Frontal Gyrus | 30% | 44.3 | 28.7 | 19.5 | 0.0300 | 0.0178 | 0.3559 | Not retained after BH |
| All participants | Glx | Cortisone | Sex interaction — within-subject component | 4 | 83 | 10,902 | Left | Frontal Pole | 83% | -42.1 | 43.9 | 14.4 | 0.0310 | 0.0178 | 0.3559 | Not retained after BH |
| All participants | Inositol | Cortisol | Between-subject effect | 1 | 5,374 | 705,900 | Left | Superior Frontal Gyrus | 49% | -16.7 | 33.8 | 55.1 | 0.0080 | 0.0080 | 0.1180 | Not retained after BH |
| All participants | Inositol | Cortisone | Sex interaction — within-subject component | 1 | 6,389 | 839,225 | Left | Lateral Occipital Cortex, superior division | 44% | -42.1 | -62.8 | 44.9 | 0.0014 | 0.0014 | 0.0280 | Retained after BH |
| All participants | Inositol | Cortisone / Cortisol ratio | Between-subject effect | 1 | 2,012 | 264,286 | Left | Left Cerebral White Matter | 100% | -21.7 | -22.2 | 34.8 | 0.0184 | 0.0184 | 0.1226 | Not retained after BH |
| All participants | Inositol | Cortisone / Cortisol ratio | Sex interaction — within-subject component | 1 | 72 | 9,458 | Right | Precentral Gyrus | 28% | 49.4 | -1.8 | 34.8 | 0.0382 | 0.0382 | 0.3819 | Not retained after BH |
| All participants | Inositol | Cortisone / Cortisol ratio | Sex interaction — within-subject component | 2 | 49 | 6,436 | Left | Frontal Pole | 71% | -47.2 | 49.0 | -5.9 | 0.0392 | 0.0382 | 0.3819 | Not retained after BH |
| All participants | Inositol | Cortisone / Cortisol ratio | Sex interaction — within-subject component | 3 | 36 | 4,729 | Left | Precentral Gyrus | 30% | -31.9 | -12.0 | 50.0 | 0.0430 | 0.0382 | 0.3819 | Not retained after BH |
| All participants | Inositol | Cortisone / Cortisol ratio | Sex interaction — within-subject component | 4 | 32 | 4,203 | Left | Superior Frontal Gyrus | 54% | -11.6 | -1.8 | 70.3 | 0.0396 | 0.0382 | 0.3819 | Not retained after BH |
| All participants | Inositol | Cortisone / Cortisol ratio | Sex interaction — within-subject component | 5 | 19 | 2,496 | Right | Right Cerebral White Matter | 100% | 13.8 | 33.8 | 9.3 | 0.0402 | 0.0382 | 0.3819 | Not retained after BH |
| All participants | Inositol | Cortisone / Cortisol ratio | Sex interaction — within-subject component | 6 | 11 | 1,445 | Left | Postcentral Gyrus | 34% | -52.2 | -32.3 | 55.1 | 0.0416 | 0.0382 | 0.3819 | Not retained after BH |

| Population | Metabolite | Hormone / predictor | Effect | Cluster | k (voxels) | Volume (mm$^3$) | Hemisphere | Atlas peak label | Atlas prob. | Peak x (MNI mm) | Peak y (MNI mm) | Peak z (MNI mm) | Cluster peak p-FWE | Analysis min p-FWE | BH q-value | BH status |
|---|---|---|---|---|---|---|---|---|---|---|---|---|---|---|---|---|
| All participants | Inositol | DHEA-S | Between-subject effect | 1 | 3,079 | 404,441 | Left | Superior Parietal Lobule | 29% | -21.7 | -52.7 | 55.1 | 0.0118 | 0.0118 | 0.1180 | Not retained after BH |
| All participants | tNAA | Cortisol / 11-deoxycortisol ratio | Between-subject effect | 1 | 895 | 117,562 | Left | Lateral Occipital Cortex, superior division | 23% | -37.0 | -67.9 | 29.7 | 0.0282 | 0.0282 | 0.5639 | Not retained after BH |
| All participants | tNAA | Cortisol / 11-deoxycortisol ratio | Sex interaction — between-subject component | 1 | 18 | 2,364 | Left | Frontal Pole | 6% | -11.6 | 54.1 | 19.5 | 0.0434 | 0.0434 | 0.3439 | Not retained after BH |
| All participants | tNAA | Cortisone | Sex interaction — within-subject component | 1 | 2,793 | 366,874 | Left | Postcentral Gyrus | 35% | -42.1 | -22.2 | 55.1 | 0.0168 | 0.0168 | 0.3359 | Not retained after BH |
| All participants | Creatine | Age | Within-subject effect | 1 | 450 | 59,110 | Right | Superior Parietal Lobule | 29% | 8.8 | -52.7 | 70.3 | 0.0226 | 0.0226 | 0.0678 | Not retained after BH |
| All participants | Creatine | Age | Within-subject effect | 2 | 292 | 38,356 | Left | Precentral Gyrus | 9% | -57.3 | 8.3 | 39.8 | 0.0242 | 0.0226 | 0.0678 | Not retained after BH |
| All participants | Creatine | Age | Within-subject effect | 3 | 107 | 14,055 | Left | Frontal Pole | 65% | -11.6 | 64.3 | 14.4 | 0.0314 | 0.0226 | 0.0678 | Not retained after BH |
| All participants | Creatine | Age | Within-subject effect | 4 | 17 | 2,233 | Left | Frontal Pole | 78% | -37.0 | 54.1 | 4.3 | 0.0428 | 0.0226 | 0.0678 | Not retained after BH |
| All participants | Creatine | Age | Within-subject effect | 5 | 16 | 2,102 | Left | Inferior Frontal Gyrus, pars opercularis | 74% | -57.3 | 13.4 | 14.4 | 0.0382 | 0.0226 | 0.0678 | Not retained after BH |
| All participants | Creatine | Age | Between-subject effect | 1 | 772 | 101,406 | Left | Paracingulate Gyrus | 62% | -6.5 | 18.5 | 39.8 | 0.0100 | 0.0100 | 0.0600 | Not retained after BH |
| All participants | Creatine | Age | Between-subject effect | 2 | 125 | 16,419 | Right | Lateral Occipital Cortex, inferior division | 16% | 34.2 | -73.0 | 9.3 | 0.0266 | 0.0100 | 0.0600 | Not retained after BH |
| All participants | Creatine | Age | Between-subject effect | 3 | 60 | 7,881 | Left | Occipital Pole | 62% | -16.7 | -98.4 | -11.0 | 0.0390 | 0.0100 | 0.0600 | Not retained after BH |
| All participants | Glx | Age | Between-subject effect | 1 | 4,366 | 573,495 | Left | Middle Frontal Gyrus | 32% | -26.8 | 18.5 | 44.9 | 0.0014 | 0.0014 | 0.0084 | Retained after BH |
| All participants | Inositol | Age | Within-subject effect | 1 | 1,294 | 169,973 | Left | Postcentral Gyrus | 51% | -62.4 | -6.9 | 34.8 | 0.0260 | 0.0260 | 0.1560 | Not retained after BH |
| All participants | Inositol | Age | Within-subject effect | 2 | 70 | 9,195 | Left | Parahippocampal Gyrus, posterior division | 18% | -31.9 | -27.2 | -16.1 | 0.0360 | 0.0260 | 0.1560 | Not retained after BH |
| All participants | Inositol | Age | Within-subject effect | 3 | 15 | 1,970 | Left | Left Putamen | 100% | -21.7 | 8.3 | -5.9 | 0.0422 | 0.0260 | 0.1560 | Not retained after BH |
| All participants | tNAA | Age | Within-subject effect | 1 | 1,660 | 218,049 | Right | Inferior Temporal Gyrus, temporooccipital part | 50% | 49.4 | -57.7 | -16.1 | 0.0146 | 0.0146 | 0.0438 | Retained after BH |
| All participants | tNAA | Age | Within-subject effect | 2 | 18 | 2,364 | Left | Precentral Gyrus | 50% | -1.4 | -27.2 | 50.0 | 0.0440 | 0.0146 | 0.0438 | Retained after BH |
| All participants | tNAA | Age | Main effect of sex | 1 | 3,172 | 416,657 | Left | Planum Temporale | 33% | -62.4 | -17.1 | 9.3 | 0.0144 | 0.0144 | 0.0438 | Retained after BH |
| Males | Choline | Testosterone | Within-subject effect | 1 | 13 | 1,708 | Right | Superior Temporal Gyrus, posterior division | 22% | 49.4 | -32.3 | 4.3 | 0.0232 | 0.0206 | 0.0824 | Not retained after BH |
| Males | Choline | Testosterone | Within-subject effect | 2 | 12 | 1,576 | Right | Intracalcarine Cortex | 43% | 3.7 | -78.1 | 4.3 | 0.0206 | 0.0206 | 0.0824 | Not retained after BH |
| Males | Glx | Testosterone | Within-subject effect | 1 | 2,497 | 327,993 | Left | Left Cerebral White Matter | 100% | -21.7 | 28.7 | 24.6 | 0.0018 | 0.0018 | 0.0072 | Retained after BH |
| Males | tNAA | Testosterone | Within-subject effect | 1 | 1,823 | 239,460 | Right | Lateral Occipital Cortex, superior division | 48% | 44.3 | -83.2 | 24.6 | 0.0092 | 0.0092 | 0.0368 | Retained after BH |
| Females | Choline | Estradiol and progesterone | Estradiol — between-subject effect | 1 | 786 | 103,245 | Right | Middle Frontal Gyrus | 19% | 29.1 | -1.8 | 50.0 | 0.0050 | 0.0050 | 0.0400 | Retained after BH |
| Females | Choline | Estradiol and progesterone | Estradiol — between-subject effect | 2 | 135 | 17,733 | Right | Lateral Occipital Cortex, superior division | 71% | 29.1 | -73.0 | 44.9 | 0.0282 | 0.0050 | 0.0400 | Retained after BH |
| Females | Choline | Estradiol and progesterone | Estradiol — between-subject effect | 3 | 14 | 1,839 | Right | Frontal Pole | 72% | 49.4 | 38.8 | 14.4 | 0.0386 | 0.0050 | 0.0400 | Retained after BH |
| Females | Creatine | Estradiol and progesterone | Progesterone — within-subject effect | 1 | 11 | 1,445 | Left | Left Cerebral White Matter | 87% | -1.4 | -1.8 | 9.3 | 0.0384 | 0.0328 | 0.1312 | Not retained after BH |
| Females | Creatine | Estradiol and progesterone | Estradiol — within-subject effect | 1 | 95 | 12,479 | Left | Planum Polare | 1% | -37.0 | -17.1 | -11.0 | 0.0134 | 0.0134 | 0.1072 | Not retained after BH |
| Females | Glx | Estradiol and progesterone | Estradiol — between-subject effect | 1 | 156 | 20,491 | Left | Superior Parietal Lobule | 40% | -42.1 | -42.5 | 55.1 | 0.0340 | 0.0232 | 0.1856 | Not retained after BH |
| Females | Glx | Estradiol and progesterone | Estradiol — between-subject effect | 2 | 142 | 18,652 | Right | Frontal Pole | 62% | 18.9 | 54.1 | 24.6 | 0.0232 | 0.0232 | 0.1856 | Not retained after BH |
| Females | Glx | Estradiol and progesterone | Estradiol — between-subject effect | 3 | 27 | 3,547 | Right | Parahippocampal Gyrus, posterior division | 66% | 18.9 | -27.2 | -21.2 | 0.0410 | 0.0232 | 0.1856 | Not retained after BH |
| Females | Glx | Estradiol and progesterone | Estradiol — between-subject effect | 4 | 14 | 1,839 | Left | Middle Frontal Gyrus | 56% | -47.2 | 8.3 | 44.9 | 0.0470 | 0.0232 | 0.1856 | Not retained after BH |
| Females | tNAA | Estradiol and progesterone | Estradiol — between-subject effect | 1 | 676 | 88,796 | Left | Postcentral Gyrus | 59% | -57.3 | -12.0 | 39.8 | 0.0174 | 0.0174 | 0.1392 | Not retained after BH |
